# Shifts in marine invertebrate bacterial assemblages associated with tissue necrosis during a heatwave

**DOI:** 10.1101/2021.01.25.428091

**Authors:** Esther Rubio-Portillo, Alfonso A. Ramos Esplá, Josefa Antón

## Abstract

Marine heatwaves (MHWs) are periods of extremely high seawater temperature that affect marine ecosystems in several ways. Anthozoans (corals and gorgonians) and Porifera (sponges) are usually among the taxa most affected by MHWs. Both are holobiont entities that form complex interactions with a wide range of microbes, which are an essential part of these organisms and play key roles in their health status. Here, we determine microbial community changes suffered in two corals (*Cladocora caespitosa* and *Oculina patagonica*), one gorgonian (*Leptogorgia sarmentosa*), and one sponge (*Sarcotragus fasciculatus*) during the 2015 MHW. The microbial communities were different among hosts and displayed shifts related to host health status, with a higher abundance in necrosed tissues of *Ruegeria* species or of potential pathogens like *Vibrio.* We also carry out a meta-analysis using 93 publicly accessible 16S rRNA gene libraries from *O. patagonica*, *C. caespitosa* and *L. sarmentosa* to establish a Mediterranean core microbiome in these species. We have identified one *Ruegeria* OTU that maintained a stable and consistent association with these species, which was also related with tissue necrosis in their hosts. Therefore, *Ruegeria* sp. could play an important and still underexplored role in the health status of its hosts.

## Introduction

Marine heatwaves (MHWs) are periods of extremely high seawater temperature that persist for days to months and can extend up to thousands of kilometers (Frölicher and Laufkötter, 2018). Some of the recently observed marine heatwaves revealed the high vulnerability of marine ecosystems, which can be affected in several ways, such as by decreasing productivity, altering food web dynamics, shifting species distribution, and reducing abundance (Hughes et al., 2003; Hoegh-Guldberg and Bruno, 2010). MHWs, which will probably intensify with anthropogenic climate change (Frölicher and Laufkötter, 2018), are related to mass mortality events and disease outbreaks in marine species that severely threaten the structure and functioning of ecosystems and disrupt the provision of ecological goods and services in coming decades (Smale et al., 2019). The most recently observed marine heatwave with global ecological implications was recorded in 2015/16, when unusually high ocean temperatures associated with one of the strongest El Niño events on record triggered unprecedented coral bleaching and marine invertebrate mortality worldwide (Rubio-Portillo et al., 2016a; Ampou et al., 2017; Oliver et al., 2017; Turicca et al., 2018).

Anthozoa (Scleractinians and Octocorals) and Porifera are important members of the benthic community. These taxa provide structural complexity to ecosystems and thereby refuge and habitats to other fauna and are the taxa most affected by MHWs (Cerrano et al., 2000; Garrabou et al., 2009). Like all multicellular organisms, marine benthic invertebrates (encompassing Anthozoa and Porifera) are holobiont entities, forming complex interactions with a wide range of microbes, including dinoflagellates, fungi, bacteria, archaea, and viruses (Knowlton and Rohwer, 2003). These microbial symbionts play active roles in holobiont health (e.g., nutrient supply and protection against pathogens) as well as the adaptive response of the host to environmental changes (reviewed in Bourne et al., 2016 and Pita et al., 2018).

Changes in the environment may severely disturb host-microbe interactions and thus lead to dysbiosis (microbial imbalance on or inside the host) and/or disease development (Harvell et al., 2007; Miller and Richardson, 2014; Sweet et al., 2015). Therefore, the evaluation of the shifts in microbiota as a result of MHWs may be employed as “early” bio-indicators of both environmental changes and host disease. However, few studies have investigated the microbiota of marine invertebrates other than corals during warming events. Microbial community association with marine invertebrates is dynamic and includes a ubiquitous core microbiome, which is defined as stable and consistent components across complex microbial assemblages from similar habitats (see review by Sweet and Bulling, 2017). These core members play key roles due to their ability to maintain microbial associations’ stability under environmental changes through competition for nutrients and/or space with invasive microbes, as well as by production of antibiotics (Ritchie, 2006; Krediet et al., 2013). Along with core members, there is a second associated microbial fraction that is more influenced by the local environmental conditions and a third highly variable component dependent on the processes occurring at the spatial and temporal scales (Reveillaud et al., 2014; Ainswoth et al., 2015; Hernandez-Agreda et al., 2018). Given the likely critical contribution of microbes to invertebrate holobiont adaptation to environmental changes, shifts in marine invertebrates’ microbial assemblages could be ideal indicators for host heat stress.

In the last 20 years, Mediterranean marine invertebrates have suffered an increase of disease outbreaks due to warming events (Cerrano et al., 2000; Garrabou et al., 2009; Stabili et al., 2012; Jiménez et al., 2016). Specifically, during the 2015 MHW in the Marine Protected Area of Tabarca, more than 40% of the population of the sponge *Sarcotragus fascicualatus*, the corals *Cladocora caespitosa* and *Oculina patagonica*, as well as the gorgonian *Leptogorgia sarmentosa* showed tissue necrosis signs as a consequence of the increase of seawater temperature (Rubio-Portillo et al., 2016a). Thus, the main goal of this study was to assess the effect of marine heat waves on microbial assemblages associated to those species to understand the influence of global warming conditions on these associations and ultimately on the health of marine invertebrates. To achieve this goal, we used Next Generation Sequencing to characterize, by means of 16S rRNA gene metabarcoding, a total of 24 marine invertebrate tissue samples from apparently healthy and necrosed colonies. We have identified potential microbial bio-indicators of marine invertebrate diseases, such as the increase of *Ruegeria* and *Vibrio* genera and a decrease of putative symbionts like *Pseudovibrio* or *Endozoicomonas* in necrosed tissues of corals and gorgonian, respectively. Moreover, a meta-analysis using 93 publicly accessible 16S rRNA gene libraries from *O. patagonica*, *C. caespitosa* and *L. sarmentosa* was carried out to establish a Mediterranean core microbiome in these species. We determined a stable and consistent association between a *Ruegeria* OTU and geographically and phylogenetic distinct Mediterranean Anthozoans, which could play an important role in the host health status. Further, our results suggest that the composition of the core microbiome depends on the geographical area considered in the analysis, confirming the existence of a local core microbiome that depends on the surrounding environment.

## Material and methods

### Sample collection

Water and invertebrate samples were collected on 28 September 2015 in two sampling locations in the Marine Protected Area of Tabarca. The gorgonian *L. sarmentosa* was collected at 25m depth (38°09′35″ N, 00°27′55″E), while the coral *O. patagonica* and the sponge *S. fasciculatus* were collected at 5m depth (38°09′59″ N, 00°28′56″E, Spain). For each of the three sampled marine invertebrates, a total of six (3 healthy and 3 necrosed) independent specimens were taken. In addition, two water samples were taken from each sampling location (Table 1). The health status of the invertebrates and the environmental parameters are described in Rubio-Portillo et al., (2016a).

All samples were taken during the heat wave recorded in September 2015. During this MHW, water temperature was 2 °C higher compared with the preceding 9 years, and persisted for approximately 6 weeks, reaching a maximum of 28.23°C (Rubio-Portillo et al., 2016a). Marine invertebrate samples were removed by SCUBA diving using a hammer and chisel and placed in plastic bags under water. Two water samples were taken from each depth using sterilized bottles. All samples were transported to the laboratory in a cooler within the next 2 hours. In the lab, marine invertebrate samples were gently washed three times with 50 ml of sterile filtered seawater (SFSW) to remove non-associated microbes and approximately 2 g (wet weight) of each sample was crushed with 5 ml SFSW using a mortar and they were allowed to settle for 15min and the supernatant (that is, crushed tissue) was removed and kept at −80 °C for further analyses.

### DNA extraction and polymerase chain reaction amplification of 16S rRNA genes

DNA was extracted from crushed tissue using the UltraClean Soil DNA Kit (MoBio; Carlsbad, CA) following the manufacturer’s instructions for maximum yield. DNA from water samples was extracted using the DNeasy blood and tissue kit (Qiagen, Valencia, CA). The extracted genomic DNA was used for PCR amplifications of the V3-V4 region of the 16S rRNA gene by using the following universal primers: Pro341F (CCTACGGGNBGCASCAG) (Takahashi et al., 2013) and Bact805R (GACTACHVGGGTATCTAATCC) (Herlemann et al., 2011). Each PCR mixture contained 5 μl of 10x PCR reaction buffer (Invitrogen), 1.5 μl of 50 mM MgCl2, 1 μl 10 mM dNTP mixture, 1 μl of 100 μM of each primer, 1 unit of Taq polymerase, 3 μl of BSA (New England BioLabs), sterile MilliQ water up to 50 μl and 10 ng of DNA. Negative controls (with no template DNA) were included to assess potential contamination of reagents. The amplification products were purified with the GeneJET PCR purification kit (Fermentas, EU), quantified using the Qubit Kit (Invitrogen), and the quality (integrity and presence of a unique band) was confirmed by 1% agarose gel electrophoresis. Sequencing was performed using Illumina Mi-seq Nextera 2×300 bp paired-end run (at Fundació per al Foment de la Investigació Sanitària i Biomédica, FISABIO, Valencia).

### Illumina high-throughput 16S rRNA gene sequence analysis

Paired-end MiSeq sequences of the 22 samples were deposited in the NCBI Sequence Read Archive (SRA) database. Data from the water samples as well as *O. patagonica, L. sarmentosa* and *S. fasciculatus* were deposited under BioProject PRJNA615777. For comparative purposes, sequences from the coral *C. caespitosa* (BioProject PRJNA407809) were also included in the analysis. These *C. caespitosa* samples were taken at 5 m depth location in the same sampling campaign than the samples listed in Table 1 and were used for a previous biogeography study (Rubio-Portillo et al., 2018). The QIIME 1.8.0 pipeline (Caporaso et al., 2010) was used for data processing. Operational taxonomic units (OTUs) were defined at the level of 99% similarity, close to the threshold used to distinguish species (98.7% similarity in the whole 16S rRNA gene), (Stackebrandt and Ebers, 2006), followed by taxonomy using UCLUST algorithm (Edgar, 2010) with the SILVA reference database (version 132). OTUs classified as chloroplast or mitochondria were removed from the dataset. Due to the large difference in library size among samples, the OTU table was rarefied to 11,594 reads (the lowest number of the post-assembly and filtered sequences in a sample, Table S1) for comparisons across samples (Weiss et al., 2015).

### Analysis of alpha-diversity

Prokaryotic α-diversity was estimated in QIIME prior to deleting singletons and OTUs with less than 0.05% of abundance. Specifically, diversity was characterized using the Shannon diversity index and OTU richness. Differences in alpha diversity index were statistically evaluated using ANOVA analysis in R with the ‘vegan’ package (Oksanen, 2011). Prior to ANOVA, homogeneity of variance was confirmed with Cochran’s test (Cochran, 1951) and data was analyzed according to a two-factor model, where the main factors were host (i.e. marine invertebrate species) and health status. If the variances were significantly different at p = 0.05, post-hoc analyses were conducted using Student–Newman–Keuls (SNK) multiple comparisons (Underwood, 1997).

### Analysis of beta-diversity

Prior to analysis of β-diversity, singletons and OTUs with less than 0.05% of abundance were removed from the dataset. For β-diversity analysis, we used QIIME software and clustering based on the weighted UniFrac (Lozupone and Knight, 2005). To visualize microbiota similarity, we generated principal coordinate analysis (PCoA) plots from the distance matrices. Multivariate analyses were used to compare composition of microbial communities associated with the different marine invertebrate species. Similarity percentage (SIMPER) was used to identify OTUs that could be potentially responsible for these differences.

### Core microbiome meta-analysis

In order to identify the core microbiome in the studied area, phylotypes consistently present in 100% of the samples (both healthy and necrosed) from each holobiont were considered. We used a conservative representation of the core microbiome because only six samples were recovered from each marine invertebrate during this study. In addition, to identify cosmopolitan microorganisms associated with benthic Mediterranean invertebrates, the core microbiome of *C. caespitosa, O. patagonica* and *L. sarmentosa* across the Mediterranean Sea was analyzed using the recommended cut off at 85% of the samples (Ainswoth et al., 2015; Hernandez-Agreda et al., 2016). For this purpose, a total of 93 16S rRNA libraries were analyzed (12 generated in the present work and 81 previously published (Rubio-Portillo et al., 2016b; 2018; van de Water et al., 2017; Bednarz et al., 2019); Table S2). In addition, unique and shared taxa (at the OTU level) among hosts were displayed with the “UpSet” (visualizing intersecting sets) diagram using the “R-bioconductor” package “UpSetR” (Lex et al., 2014).

## Results and discussion

To assess the effect of global warming on marine invertebrates, we investigated the differences in the microbiome of healthy and necrosed marine invertebrates during a marine heatwave in order to explore the presence of potential microbial indicators of heat stress. In addition, the core microbiome of each host was also described as well as the presence of cosmopolitan microorganisms associated with benthic Mediterranean Anthozoans.

More than 24,000 OTUs were identified in the present study but only 173 OTUs showed a relative abundance over 0.05 % and are discussed here. Invertebrate species hosted on average from 117 to 149 OTUs (149 OTUs for *O. patagonica*, 144 for *L. sarmentosa*, 134 OTUs for *C. caespitosa* and 117 for *S. fasciculatus*). Importantly, less than 20% of OTUs identified were host-specific, while about half were shared by at least three of the invertebrates studied here (Figure 1). Particularly, the three Anthozoans are the hosts that shared more OTUs among them (14.45%). Furthermore, 28 OTUs were shared among all the marine invertebrates and seawater samples (Figure 1). Therefore, it seems that the surrounding water has a great influence on the invertebrate microbiome, which is in good agreement with previous studies that showed biogeographical changes in the invertebrate microbiome (Littman et al., 2009; Pantos et al., 2015; Rubio-Portillo et al., 2018). A large proportion of sequences related to the *O. patagonica* pathogen *V. mediterranei* (Kushmaro et al., 1997;1998; Rubio-Portillo et al., 2014) was detected in seawater samples and this OTU was shared by all samples (Table S3). This fact was probably as consequences of the increasing temperature during the MHW and this could compromise benthic invertebrate health. Conversely, since vibrios have been detected in viable but not culturable state in coral tissue during cold seasons (Sharon and Rosenberg, 2010; Rubio-Portillo et al., 2016b), invertebrates could act as a pathogen reservoir, from which they could be dispersed into the surrounding water.

**Figure 1.**
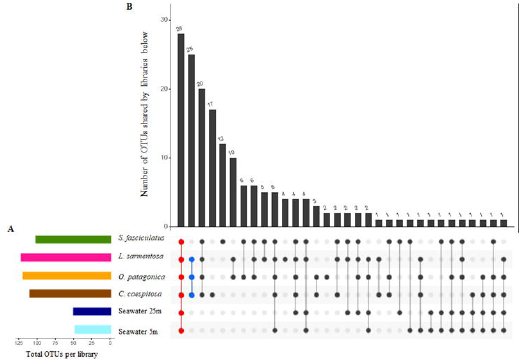
Upset plot showing the relationship of OTUs identified in all marine invertebrates and seawater samples analyzed in this study. A) Graph of the OTUs average (X axis) in each sample (Y axis). B) Intersection of sets of OTUs in each sample. The number of OTUs in each set appears above the column, while the sample shared are indicated in the graph below the column by a point, with the samples on the left. Intersection in red represent OTUs shared by all hosts and seawater samples and intersection in blue OTUs shared by the three Antozoan.

The Shannon diversity index ranged from 2 to 5 in the marine invertebrates studied here, consistent with previous studies (Rubio-Portillo et al., 2016b, Thomas et al., 2016; 2018; van de Water et al., 2018a). The two-way ANOVA revealed significant differences among hosts (F= 23.114, p < 0.001). Post-hoc SNK test showed that these differences were due to the highest diversity values showed by *O. patagonica* compared with the other hosts, which diversity was similar among them (Figure 2A). Similarly, OTU richness was also higher in *O. patagonica* than in the other hosts (Figure 2B; F= 14.546, p < 0.001). Principal coordinate analysis using weighted UniFrac distances (Lozupone and Knight, 2005) clearly separated samples by hosts, which were also different from seawater samples (Figure 3A and B). Bacterial microbiomes associated with the two zooxanthelate scleractian corals were similar to each other and different from the azooxanthellate gorgonian *L. sarmentosa* microbiome (Fig. 3A and 3B). For instance, *Endozoicomonas* genus, a common coral symbiont (Bourne et al., 2016; Neave et al., 2016), was one of the most abundant genera in the gorgonian *L. sarmentosa* (Fig. 4B and Table 1), in good agreement with previous studies carried out in the Mediterranean Sea (Bayer et al., 2013; Rubio-Portillo et al., 2016b; Rubio-Portillo et al., 2018; Van de Water et al., 2017). However, intriguingly, this genus was absent from the corals studied here. In addition to differences among Anthozoans, differences between the two coral species were also observed. SIMPER analysis showed that *Maritimimonas* was a characteristic genera of *C. caespitosa*, while *Pseudovibrio* genus was significantly enriched in *O. patagonica* (Table S4). Likewise, SIMPER analysis revealed that sequences corresponding to uncultured genera of *Acidobacteria* and *Dadabacteria* were sponge-specific (Table S4). Therefore, although surrounding water had a great influence on the invertebrate microbiome, the microbial composition was different for the different hosts and specific symbionts were detected in each host.

**Figure 2.**
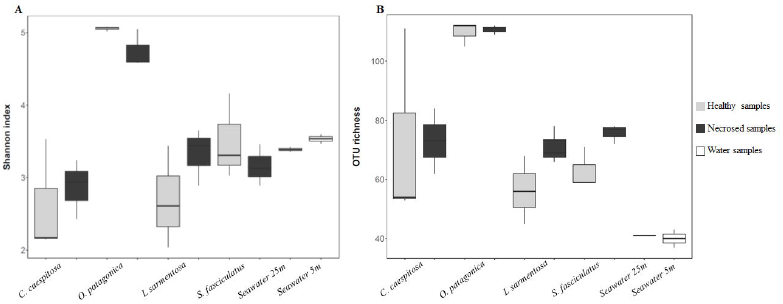
(A) Shannon diversity index and (B) OTU richness obtained for host-associated and surrounding water microbiomes based on 16S rRNA gene diversity.

**Figure 3.**
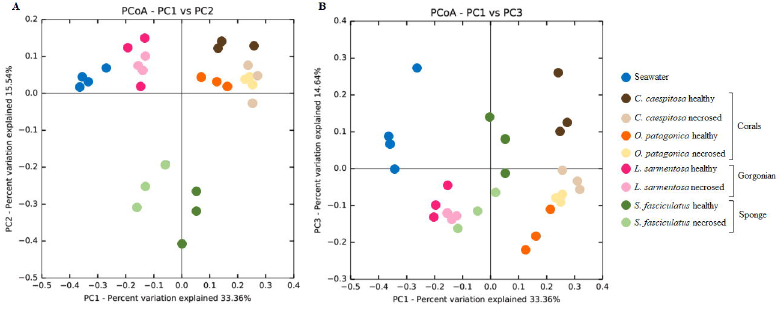
Principle coordinate analysis (PcoA) 2D plot based on microbial communities associated with *Cladocora caespitosa, Oculina patagonica, Leptogorgia sarmentosa* and *Sarcotragus fasciculatus* tissues clustered using coordinated analysis of the weighed UniFrac distance matrix. A) The x- and y-axes are indicated by the first and second coordinates, respectively, and the values in parentheses show the percentages of the community variation explained. B) The x- and y-axes are indicated by the first and third coordinates, respectively, and the values in parentheses show the percentages of the community variation explained

**Figure 4.**
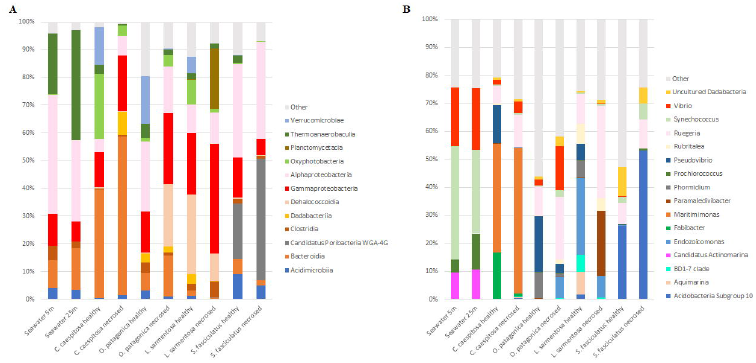
Overview of the composition of the microbiome composition and microbial community changes related to tissue necrosis signs associated with *Cladocora caespitosa, Oculina patagonica, Leptogorgia sarmentosa* and *Sarcotragus fasciculatus* at (A) class and (B) genus level. For full taxonomic information refer to Supplementary Supplementary Data 1.

## Microbiota shifts related to host health status

As shown in figures 2 and 3, although there were no detectable differences in terms of diversity indexes, microbial composition changed depending on health status. Thus, both Shannon index and OTU richness did not show significant changes among healthy and necrosed samples in either host species (Fig. 2A an 2B; F= 0.796, p =0.5141). However, PERMANOVA analysis (R^2^= 0.651, p < 0.005) as well as principal coordinate analysis using weighted UniFrac distances showed that microbial composition differed depending on health status (Fig. 3A and 3B). SIMPER analysis was carried out in order to detect the OTUs primarily responsible for these differences (Table S5). For the analyzed Anthozoans, this analysis unveiled a common pattern in the necrosed tissues compared with the healthy ones, with a decrease of potential symbionts like *Pseudovibrio, Fabibacter* or *Endozocicomas* genera and an increase of opportunistic and *Vibrio* species, together with the increase of some species whose role is still not clear, like *Ruegeria* spp. (Fig. 4B, Table S6 and Table S4).

*Pseudovibrio* species, which play a key role in coral health by inhibiting pathogens′ growth (Nissimov et al., 2009; Rypien et al., 2010), were more abundant in healthy corals than in necrosed ones. Thus, the *Pseudovibrio*-dominated community changes to a community dominated by potential pathogens in *O. patagonica* necrosed samples. The same pattern was also observed in samples collected in the same studied area in 2011 (Rubio-Portillo et al., 2016b). Therefore, this genus appears to be a vital member of the *O. patagonica* holobiont and its abundance could be an indicator of host health. *Pseudovibrio* OTUs detected in the two coral species studied here were different, which suggests that different coral species could select different symbionts in the same environment. In addition to *Pseudovibrio, Fabibacter* showed higher abundance in healthy specimens of *C. caespitosa* and it could also play a key role in host health. *Fabibacter* species has been previously reported associated to other coral species (Sunagawa *et al*., 2009; De castro et al., 2010), but their role remains unknown. For the gorgonian *L. sarmentosa*, tissue damage was associated with a decrease of species commonly associated with gorgonians like *Endozoicomonas* (Figure 4B and Table S4). *Endozoicomonas* spp. are one of the main constituents of octocoral microbial assemblages in the Mediterranean Sea (that can make up to over 96% of the community) and a decrease in its abundance has been correlated to environmental stress (reviewed in van de Water et al., 2018b). This is one of the key findings of this work and highlights the importance of *Pseudovibrio*, and probably *Fabibacter*, together with Endozoicomonas genera in Mediterranean Anthozoans. These microbes could serve as potential indicators of compromised health status in Mediterranean corals and gorgonians, respectively.

Intriguingly, although the relative abundance of *Vibrio* spp. increased in *O. patagonica* and slightly in *C. caespitosa* necrosed samples (Figure 4B and Table S6), SIMPER analysis did not detect any specific *Vibrio* OTU as primarily responsible for these differences (Table S4). For example, the coral pathogens *Vibrio mediterranei* and *Vibrio coralliilyticus* (OTUs 163 and 165, respectively) were detected in necrotic tissues but also in apparently healthy specimens at the same location (Table S5). This result suggests that probably the strains detected in healthy and necrosed samples could be different and not all of them pathogenic. Indeed, *V. mediterranei* strains similar to the type strain AK-1, the causative agent of mass bleaching events in *O. patagonica*, were mainly isolated from the necrosed specimens of *O. patagonica* (Rubio-Portillo et al., 2016). Along with the increase of *Vibrio* species, we have detected a consistent increase of *Ruegeria* sp. SOEmb9 OTU119 in necrosed tissues of all Anthozoans studied here (Table S5). Previous studies have shown that the presence of *Ruegeria* species is correlated with the presence of *Vibrio* pathogens in coral tissues (Rosado et al., 2019), as well as with different signs of disease, such as Black Band Disease in the Caribbean Sea (Sekar et al., 2008), Yellow Band Disease in the Red Sea (Apprill et al., 2013) or White Patch Syndrome in the Indian Ocean (Séré et al., 2013). Furthermore, *Ruegeria* genus, belonging to the *Roseobacter* group, displays high chemotactic attraction towards dimethylsulfoniopropionate (DMSP) (Miller et al., 2004), which is a compound found in heat-stressed zooxanthelate corals (Raina et al., 2013), This behavior could explain the increase of *Ruegeria* sp. in zooxanthelate coral necrotic tissues during this mortality event, probably due to the increase of DMSP as a result of the increase of sea water temperature during the heatwave. However, there are alternative explanations since some studies showed that *Ruegeria* spp. have an important role protecting corals against pathogenic *Vibrio* species by inhibiting their growth (Miura et al., 2018; Rosado et al., 2019). Therefore, further experimental evidence would be necessary in order to elucidate the role of this genus in mortality events in marine azooxanthellate invertebrates.

Overall, our results show that together with *Vibrio* coral pathogens, other specific indicators should be used to assess marine invertebrates’ heat stress and *Ruegeria* is likely a good candidate.

In the sponge *S. fasciculatus*, the changes related to health status were less evident compared to Anthozoan species. Sponge microbiome has been described to be dominated by *Proteobacteria* with *Chloroflexi, Cyanobacteria* and *Crenarchaeota* occasionally reaching high relative abundances (Thomas et al., 2016). In the current study, almost the same phyla were present in healthy and necrosed sponge microbiomes, which were dominated by *Proteobacteria* and *Poribacteria*, although some differences could be detected at genus level. *Acidobacteria* Subgroup 10 became dominant in necrosed samples (Figure 4B and Table S6) compared to healthy samples. This increase was due to OTU8 (Table S5), which was closely related to an uncultured *Acidobacteria* clone (FJ269280.1) isolated from the sponge *Xestospongia testudinaria* in Indonesia (Montalvo and Hill, 2011). This finding suggests that this OTU could be shared by taxonomic and geographically distant sponge hosts and could be a generalist symbiont within the core sponge microbiome. An increase of *Synechoccus* genus was also detected in necrosed sponges (Figure 4B and Table S6). However, one of the main OTUs responsible for the differences among healthy and necrosed samples in *S. fasciculatus* was OTU57 related to *Candidatus* Synechococcus spongiarum (Slaby and Hentschel, 2017). This OTU was a characteristic of healthy samples, where it was 3-fold more abundant than in necrosed ones (Table S5). *Candidatus* S. spongiarum was previously reported as one of the most common symbionts in this sponge and its abundance was related to the increase of seawater temperature (Erwin et al., 2012). Similar shifts in *S. fasciculatus* associated bacteria community composition have been reported previously during the 2010 summer disease episode in the Mediterranean Sea with higher abundances of *Acidobacteria* and lower abundances of *Candidatus* S. spongiarum (Blanquer et al., 2016), although the diseases signs observed by these authors (small white spots) were different to the tissue necrosis reported in the current study. Thus, it seems that an increase of *Acidobacteria* Subgroup 10 and a decrease of *S. spongiarum* could be indicators of heat stress in *S. fasciculatus*, although more studies are necessary in order to understand their role in the sponge diseases development.

## Core microbiome of Anthozoans in the Mediterranean Sea

Previous studies (van de Water et al., 2017; 2018a) have demonstrated that the microbiome of Mediterranean Anthozoans largely depends on their location and could influence their hosts’ adaptation to new environmental conditions. Therefore, in order to ascertain the core microbiome associated with each of the three Anthozoans studied here (*C. caespitosa, O. patagonica* and *L. sarmentosa*) throughout the Mediterranean Sea, we have analyzed a total of 98 16S rRNA libraries from different Mediterranean locations (Table S1), including a total of 143,281 OTUs. The analysis indicated that the core microbiome of these three Anthozoans species in the studied area was composed of 43 OTUs in *C. caespitosa*, 45 in *O. patagonica* and 43 in *L. sarmentosa*, while this core microbiome throughout the Mediterranean Sea was reduced to 4 OTUs in *O. patagonica* (3 *Ruegeria* OTUs and 1 *Pseudovibrio* OTU), 4 in *C. caespitosa* (2 *Ruegeria* OTUs, 1 *Pseudovibrio* OTU and 1 *Vibrio owensii* OTU) and 9 in *L. sarmentosa* (5 *Endozoicomonas* OTUs, 1 *Ruegeria* OTU, 1 BD1-7 clade, 1 *Granulosicoccus* and 1 *Winogradskyella*). Thus, as expected, the increase found in the biogeographical area studied implies a decrease of the corresponding core microbiome. The presence of *Pseudovibrio* OTUs in *O. patagonica* and *C. caespitosa* and *Endozociomonas* OTUs in *L. sarmentosa* core microbiomes confirms that they are stable bacterial symbionts that are less sensitive than other members of the community to the surrounding environment and they could be good indicators of their hosts’ health. Furthermore, these results confirm that, although there is a core microbiome of Anthozoans in the Mediterranean Sea, there is also a local core, as previously observed in Mediterranean gorgonians (van de Water et al., 2017). Thus, our findings confirm that the definition of the core microbiome must be associated with the geographical area considered in the analysis.

Importantly, only one OTU, highly similar to *Ruegeria* OTU119, composed the core microbiome of these three Anthozoans. As mentioned above, *Ruegeria* OTU119 increased its abundance in unhealthy samples of all Anthozoans studied here. Although this genus has been previously related to the spread of coral diseases worldwide (Sekar et al., 2008; Sunagawa et al., 2009; Apprill et al., 2013) its role in coral microbiome is still unclear. Indeed, this genus is not only present in the core microbiome of healthy specimens *O. patagonica, C. caespitosa* and *L. sarmentosa* throughout the Mediterranean Sea, is also commonly associated with a large number of other coral species around the world (Huggett and Apprill, 2018; Rothing et al., 2020), even in larvae forms (Sharp et al., 2012; Zhou et al., 2017). It has been recently demonstrated that species belonging to this genus associated with corals show antibacterial activity against *Vibrio* coral pathogens (Miura et al., 2019) and provide essential vitamins like cobalamin (Karimi et al., 2019). Taken together, this OTU composed the core microbiome of geographically distant Anthozoans in the Mediterranean Sea and its abundance increased in necrotic tissues of their host under heat stress, highlighting the importance of this genus in marine invertebrate health during MHWs.

## Acknowledgments

The authors gratefully thank the staff of the Department of Marine Sciences and Applied Biology and the Marine Research Centre of Santa Pola (CIMAR). We also greatly appreciate the friendly cooperation of the Secretary-General for Fisheries of the Spanish Ministry of Agriculture, Food and Environment, and the Marine Reserve of Tabarca guards (particularly Felio Lozano). Authors thank Karen Neller for her professional text editing services.

